# Photostimulus intensity, rather than colour or polarization, is the key driver for stable fly visual attraction in laboratory bioassays

**DOI:** 10.64898/2026.10.02.756341

**Authors:** Emmanuel Hung, Sophie Hennig, Miele McGowan, Christina Li, Adam J. Blake, Rachel Wong, Aubrie Verleur, Hamody Sabih, Sonali Timmath, Michelle Roach, Inderpreet Lail, Gerhard Gries

## Abstract

Foraging hematophagous flies such as the stable fly, *Stomoxys calcitrans*, are shown or expected to sense multiple characteristics of visual host cues. These include relative intensity, spectral composition, and the angle and degree of polarization. Here, we investigated the role of each of these visual cue characteristics on stable fly attraction. In laboratory two-choice bioassays, we offered groups of female or male stable flies a choice between paired photostimuli that differed in their visual characteristics, recording the flies’ landing responses on stimuli as the response criterion. Bioassay flies preferentially selected lower-intensity visual stimuli even when paired stimuli markedly differed in spectral composition and polarization characteristics (a high degree of horizontal or vertical polarization). When paired stimuli had identical relative intensity, flies failed to discriminate between the stimulus that presented ultraviolet, blue, green, and red wavebands and a corresponding paired stimulus lacking one of these wavebands. Our results support the hypothesis that foraging stable flies may respond to colour and polarization characteristics of visual stimuli as modulations of perceived intensity. Acquired knowledge on the mechanisms that underlie stable fly visual attraction may inform the design of vision-based traps so that they become optimally attractive to stable flies and other biting flies.

## Introduction

Blood-feeding flies exploit host cues to orient towards and recognize suitable host animals [1,2]. When homing in on prospective hosts, foraging flies exploit visual host cues to inform directional orientation and landing decisions [1,3]. Visual host cues are so important that disrupting them may suffice to deter biting fly attacks [4,5]. It has long been known that host-foraging flies predominantly respond to dark (i.e., low-intensity) high-contrast visual stimuli [1,6]. However, the flies’ photoreceptors can sense multiple characteristics of light [7], such as the colour (i.e., spectral composition independent of intensity) and the polarization of host-reflected light [8–10], all of which may affect the flies’ host-foraging behaviour.

The fly visual system receives light primarily through paired compound eyes composed of many light-sensing units (ommatidia) [11]. Each ommatidium possesses an array of six spectrally homogenous outer photoreceptors (retinula cells R1–R6) which surround two superimposed and spectrally heterogenous inner photoreceptors (retinula cells R7 and R8), with R7 and R8 sharing a light-path [12]. All eight photoreceptors are thought to inform phototaxis [13], whereas only the outer photoreceptors are involved in dim-light vision and motion detection [14,15]. Colour vision appears to be achieved primarily by the inner photoreceptors via intra- and inter-ommatidial opponent processing [16–18]. Similarly, polarization vision may be achieved by comparing the output of inner photoreceptors whose rhabdomeres exhibit geometrically orthogonal microvillar orientation [19]. This anatomical configuration applies to several blood-feeding flies including the band-eyed brown horse fly, *Tabanus bromius*, the stable fly, and tsetse flies, *Glossina* spp. [20–22].

The potential hierarchy of, and interplay between, visual host cue characteristics (e.g., intensity, colour, polarization) that inform host-foraging behavior of blood-feeding flies are not well understood. Some innate behaviours of flies, such as phototaxis, are dependent upon the intensities of specific wavelengths [23]. For example, *Drosophila* vinegar flies exhibit a negatively intensity-dependent phototactic response to green light, but a sigmoidal intensity-response curve to ultraviolet (UV) light [24]. On the other hand, colour-opponent mechanisms may underlie object-directed colour preferences (e.g., those involved in host recognition) that are robust to changing environmental illumination [25,26]. Polarotaxis of foraging horse flies is guided by host visual cues that – compared to background surroundings – are low in reflective intensity and high in their degree of linear polarization [9,27]. In horse flies, polarization-opponent pairs of photoreceptors differ in their spectral sensitivities, rendering polarotaxis wavelength-dependent [21]. The polarization-sensitive inner photoreceptors of stable flies are likewise spectrally imbalanced [22]. Stable flies distinguish between vertical and horizontal polarization of light but only when both UV and human-visible wavelengths are presented [22]. Whether the stable flies’ preference for low-intensity objects [6] is modulated by colour and polarization characteristics is yet to be determined. Revealing the effects of, and interplay between, visual host cue characteristics on stable fly attraction may provide insight for the development of effective control tactics.

Blood-feeding stable flies are severe global pests of livestock [28]. Like other blood-feeding flies, stable flies utilize visual cues to inform their approach to prospective host animals [29,30]. Vision-based traps have long been deployed to monitor and help manage stable fly infestations [31]. However, traps are generally less attractive than host animals, and the visual cues that render traps attractive to stable flies are poorly understood [28]. A wide range of trap types have been deployed, including netted funnel traps made of blue and black fabric or plastic sheets [32–34] and adhesive traps constructed from translucent Alsynite^©^ fiberglass [35], white Coroplast^©^ [36], and other white or translucent polyethylene and poylvinyl chloride plastics [37,38]. These diverse fly-attracting trap surfaces very likely differ in their reflection and transmission of incident sunlight, making it difficult to ascertain the specific visual characteristics responsible for fly attraction.

Previous studies report that stable flies are most strongly attracted to objects with low reflective intensity [6,29] or, conversely, to objects with high blue-, UV-, or overall light-reflectance [35,39,40]. The attractiveness of Alsynite traps to stable flies is thought to be due to (*i*) their substantial reflectance of UV wavelengths (5–20%) and/or (*ii*) their polarization of transmitted light [35], but neither concept has been experimentally tested. In some stable fly studies, trap surfaces were spectrally measured [35,36]. However, in most other studies colour preferences of flies are described based on human color perception, making it difficult to interpret or replicate findings [41]. Furthermore, visual trap characteristics such as spectral intensity and polarization are often not reported and may have confounded the interpretation of results [42].

In the current study, we quantified and independently modulated the intensity, spectral composition, angle of polarization (*AoP*; the predominant angle/plane along which electric fields in a light beam oscillate) and degree of linear polarization (*DoLP*; the percentage of light waves that are oriented to the *AoP*) [43] of paired visual stimuli and tested their effects on the foraging decisions of stable flies. We tested four hypotheses (H): (H1) photostimuli of low relative intensity are most attractive to stable flies, (H2) photostimulus intensity affects fly responses more strongly than its polarization composition, (H3) UV stimuli are preferentially sought by flies, and (H4) stimulus intensities preferred by flies are wavelength-dependent.

## Materials and Methods

### Experimental flies

A new laboratory colony of stable flies was established annually using wild stable flies collected from Eagle Acres Dairy and Pumpkin Patch (Langley, BC, CA) and the Dairy Education and Research Center of the University of British Columbia (Agassiz, BC, CA). Adult flies were housed in metal mesh cages (45 × 45 × 45 cm; BioQuip®, Compton, CA, USA) and fed daily with citrated bovine blood collected from a local abattoir. Cages were held in an ER-75 walk-in growth chamber (Bio Chambers Inc., Winnipeg, MB, Canada) kept at 26 °C, 60% relative humidity, and a 14:10 h (L:D) photoperiod. All bioassay flies were 3 to 7 days old and food-deprived 20–24 h prior to bioassays.

### General bioassay design

Two-choice bioassays were run in a metal mesh cage (45 × 45 × 45 cm; BioQuip®) illuminated by an overhead metal halide lamp (MT400D/BUD/HTL-BLUE, EYE Lighting Int., Mentor, OH, USA) which was positioned 145 cm above the cage ceiling (Figure 1a), emitting a broad spectrum of UV and visible light (Figure 2a). The cage floor was lined with matte brown Kraft paper (NCR Corp., Duluth, GA, USA) to prevent unwanted light reflections, and the cage ceiling panel was replaced with eight layers of cheesecloth (Fisher Scientific, Ottawa, ON, Canada; Figure 1e) to diffuse overhead illumination. Light originating from the halide lamp also entered the cage through a side wall after being reflected by a mirror (48 × 60 cm, Lärbro, IKEA, Burlington, ON, Canada) which was covered in four layers of cheesecloth (Figure 1a). This mirror-reflected light passed through paired filter stacks (14 × 14 cm) clipped onto paired cardboard frames which, in turn, were attached to the interior face of the illuminated cage wall (Figure 1e). Filter stacks – consisting of layers of cheesecloth and black screen (#892-003, Supreme Laces, Montréal, QC, Canada) each held securely between white cardstock frames (Figures 1i, 2d–e) – modulated the transmission of mirror-reflected light according to the number of cheesecloth layers (which diffused light) and of black-screen layers (which reduced light transmission) (Table S2).

**Figure 1.**
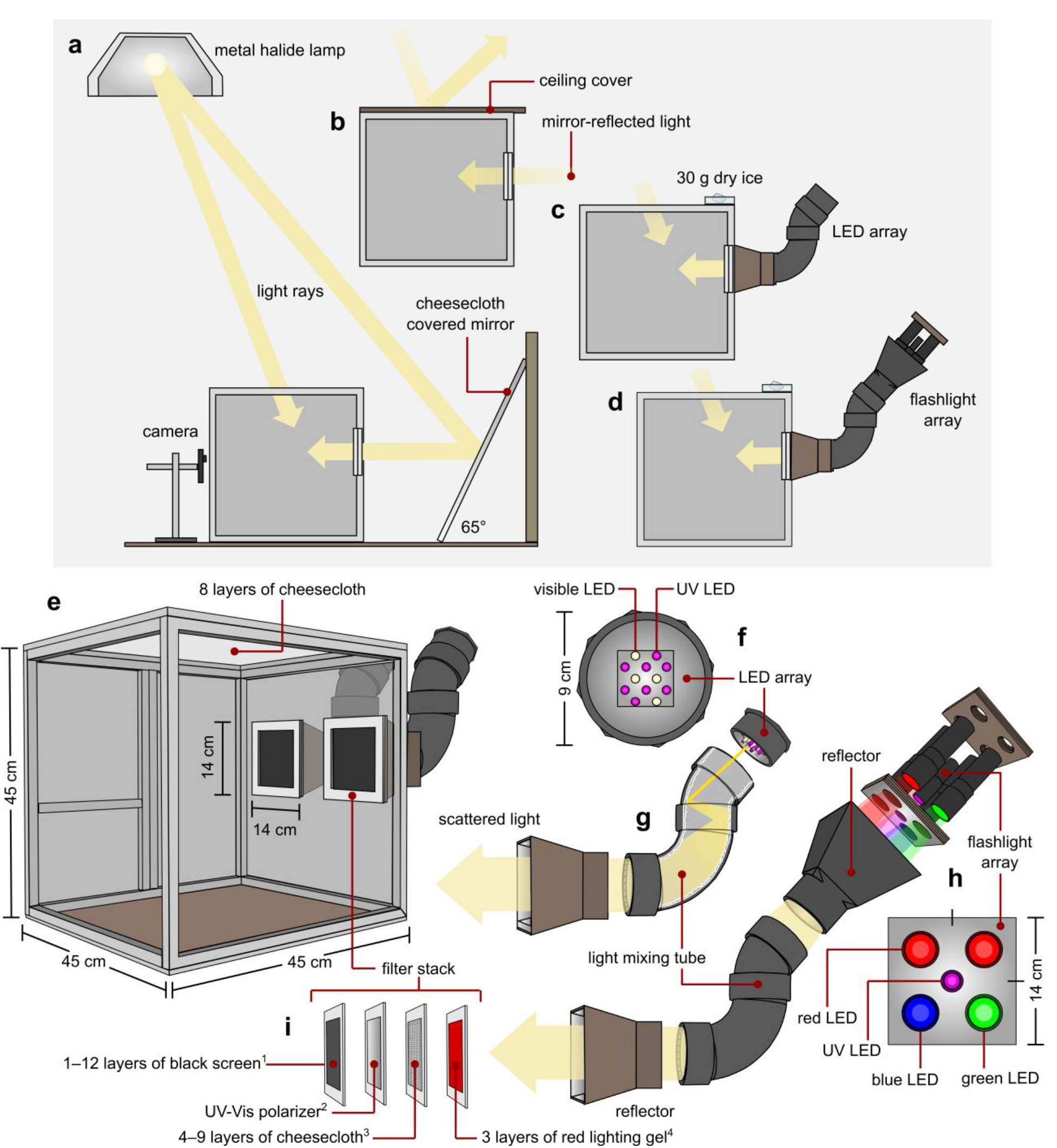
Graphical illustrations of experimental design elements and their arrangements in behavioral bioassays with stable flies. (**a–d**) Side views showing the position of (**a**) the bioassay cage relative to a mirror reflecting light from an overhead metal halide lamp, (**b**) a light-blocking cage ceiling cover used in experiments 11–20, and (**c, d**) the light-emission sources used in experiments 21–24 (**c**) and 25–52 (**d**). (**e**) Placement of paired filter stacks on the interior wall of the bioassay cage. (**f**) Frontal and side views of a light-emitting diode (LED) array, showing the arrangement of its LEDs and its placement relative to a light-mixing tube and reflector. (**g**) Interior view of a light-mixing tube with aluminum-foil lining to scatter LED-emitted light. (**h**) Frontal and side view of a flashlight array, showing the arrangement of its flashlight LEDs and its placement relative to a light mixing tube and two reflectors. (**i**) Detailed view of a filter stack showing various layers of black screen and cheese cloth, and a UV-Vis polarizer. Note: ^1^The number of black screen layers deployed modulated the light intensity and varied between paired stimuli and experiments (see Table S2); ^2^the effect of the UV-Vis polarizer on the flies’ responses was tested only in experiments 11–20; ^3^the number and position of cheesecloth layers used to diffuse light varied between paired stimuli and experiments (see Table S2); ^4^red lighting gel was applied only in experiments 38–40 and S1–S12.

**Figure 2.**
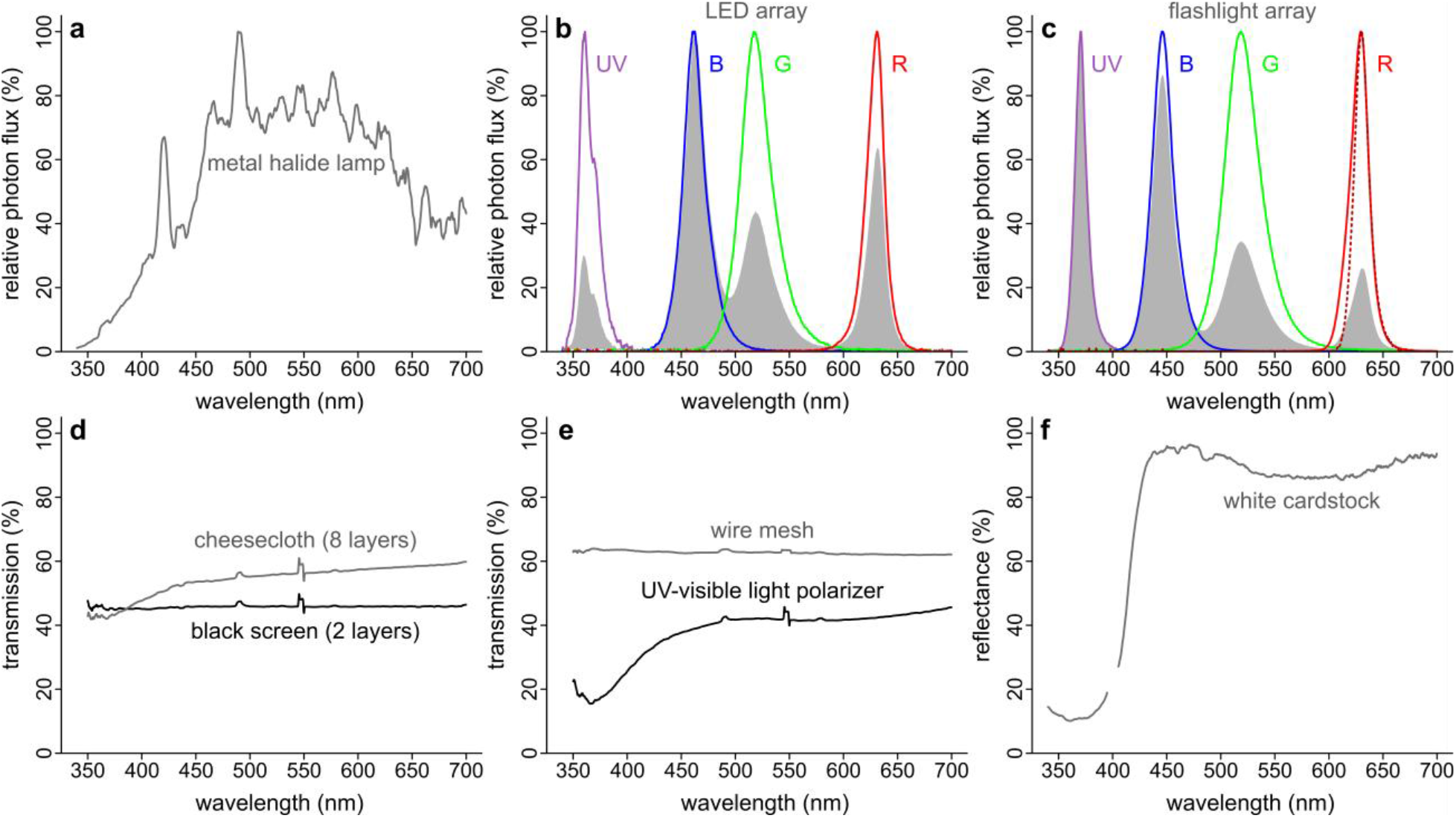
Spectral measurements of test stimuli and illuminating lights used in behavioral experiments with stable flies. (**a**) Relative photon flux of the metal halide lamp (Figure 1a); spectrum normalized by its maximum. (**b, c**) Relative photon flux of the LED array (**b**) and the flashlight array, including the red flashlight LED filtered by 6 layers of red lighting gel (dotted red line) (**c**), illuminating filter stacks via light tubes (Figure 1f–h); coloured lines show each spectrum normalized by its maximum, grey shading shows each spectrum normalized by the maximum of all spectra. (**d, e**) Transmission spectra of the wire mesh of the cage walls and the component layers of the filter stacks. (**f**) Diffuse reflectance spectra of white cardstock used in the construction of filter stacks (Figure 1i). Note: the effect of optical brighteners, which fluoresce under UV light, was eliminated in measurements above 400 nm by using a polycarbonate sheet.

At the start of each bioassay replicate, 25 food-deprived male or female stable flies were released into the cage, and their landing responses on filter stacks were recorded 5 min on a mobile phone camera (iPhone 5s or 6, Apple, Cupertino, CA, USA). The position of the treatment and the control filter stack was alternated between replicates to reduce potential side bias.

### Polarimetry

We used photo polarimetry [43,44] to measure the intensity, degree of linear polarization (*DoLP*), and the axis of polarization (*AoP*) of the photostimulus produced by each filter stack. Photos were taken using a modified Olympus E-PM1 camera (Olympus, Tokyo, Japan) (Dr. Klaus Schmitt, Weinheim, Germany, uvir.eu) [45], an ultra-broadband linear polarizing filter (68–751, Edmund Optics, USA), and both a UV/IR filter (Baader Plantarium, Mammendorf, DE) and a U-filter (Baader Plantarium) for capturing images in the human-visible (400–700 nm) and UV (330–395 nm) range, respectively. Raw images were taken of all filter stack stimuli with the polarizing filter positioned at 0°, 45°, 90° and 135°. Using a custom-created macro [46], these raw images were decoded in FIJI [46], using the DCRAW plugin, before the mean intensity, mean *DoLP*, and modal *AoP* were calculated for each of the UV (330–395 nm), blue (410–530 nm), green (455–610), and red (575–700 nm) wavebands of each filter stack stimulus.

The relative intensity, *DoLP*, and *AoP* values reported below, and in Table S1, are the mean of all four measured wavebands (Exps. 1–28) or that of a single waveband (Exps. 29–40, S1–S12) (see Table S2). Because the intensities of stimuli differed by multiple orders of magnitude across experiments, relative intensity values were normalized to the maximum intensity measured across stimuli for each set of experiments that tested a specific hypothesis (Table S2).

### Spectroscopy

We used a calibrated spectrophotometer (HR-4000, Ocean Optics Inc., Dunedin, FL, USA) to measure the emission spectra of the metal halide lamp, the LED array, and the flashlight array (Figure 2a–c), as well as the transmission of the cheesecloth, black screen, cage wire mesh, and UV-visible light polarizer through which filter stack stimuli were transmitted (Figure 2d, e). We measured the reflectance of the white cardstock used to build filter stack frames (Figure 2f) using a JAZ spectrometer (Ocean Optics) calibrated with a 99% Spectralon reflectance standard (SRS-99–010, Labsphere, NH).

### Specific experiments

#### H1: Photostimuli of low relative intensity are most attractive to flies (Exps. 1–10)

To determine the relative intensity of a photostimulus that renders it attractive to flies, flies were offered a choice between paired full-spectrum photostimuli [330–700+ nm (Figure 2a); <20% degree of linear polarization (*DoLP*); Table S2] of high relative intensity (100%, standardized control) and of lower relative intensity (9–66%; Exps. 1–5; Table S1). To determine the relative intensity differential that stable flies can discern, flies were offered a choice between a stimulus (full-spectrum, <20% *DoLP*) of low relative intensity (9%, standardized control) and of higher relative intensity (11–45%; Exps. 6–10; Table S1).

#### H2: Photostimulus intensity affects fly responses more strongly than its polarization composition (Exps. 11–20)

Drawing on reports that stable flies respond to polarized light [22], we compared the relative effects of stimulus light intensity and polarization composition on the flies’ behavioral responses. To this end, UV-visible light polarizers (Bolder Vision Optik, Boulder, CO, USA) were incorporated in filter stacks (Figure 1i), producing stimuli with high *DoLPs* (56–70%; Table S2) which – according to the polarizers’ rotation – transmitted light with an angle of polarization (*AoP*) that was predominantly horizontal (90°) or vertical (180°) (Table S2). To test whether flies retain their preference for horizontally-polarized light [22] irrespective of stimulus intensity (Exp. 11: 97–100%; Exp. 12: 58–61%; Exp. 13: 45–46%, Exp. 14: 37– 38%; Exp. 15: 28–29%), flies in each of Exps. 11–15 (Table S1) were offered a choice between paired photostimuli of near-equal relative intensity and *DoLP* but with either horizontal polarization (*AoP*: 168– 173%) or vertical polarization (*AoP*: 83–85%).

Because the flies’ preference for horizontally polarized light vanished at low(er) relative intensities (45–28%; see Results), we proceeded to determine the minimum light intensity differential between photostimuli that the flies required to preferentially respond to horizontally polarized light. To this end, flies were offered a choice between horizontally and vertically polarized photostimuli, with the horizontally polarized light stimulus set to 61% relative intensity and the vertically polarized light stimulus set to a relative intensity ranging from 28–97% (Exps. 16–20; Table S1).

Exclusively in experiments 11–20, a white cardboard cover (Figure 1b) was placed on the cage ceiling to block direct illumination of the cage wall holding the filter stacks (‘filter-stack wall’). This experimental permutation lowered the ratio of transmitted to reflected light produced by the filter stacks, thereby preserving a higher stimulus *DoLP*. In experiment 11, and in all subsequent experiments, the exterior of the two side walls perpendicular to the filter-stack wall was covered with a single layer of brown Kraft paper (NCR Corp.) to reduce cage side-dependent differences in the amount of light reflecting off the fronts of the filter stacks.

#### H3: UV stimuli are preferentially sought by flies (Exps. 21–28)

To produce paired photostimuli of specific wavebands, each of two custom-built light tubes [22] was fitted with an array of light-emitting diodes (LEDs) consisting of four RGB (red, green, blue) LEDs (N12108, BQLZR, Shenzhen, GD, China) and eight UV LEDs (EOLD-355-525, OSA Opto Light GmbH, Berlin, Germany). The LED array was housed inside a ‘light-mixing tube’ (made from ABS pipe fitting; 77 mm internal diameter) which was connected to a reflector that channeled light emissions towards a filter stack (Figure 1c,f,g). To scatter light rays, interior walls of the light-mixing tube and the reflector were lined with crumpled, re-flattened Alcan aluminum foil (Reynolds Consumer Products Canada Inc., Mississauga, ON, Canada) (Figure 1f). To test whether attraction of flies to photostimuli (<20% *DoLP*) was affected by the presence of UV (330–395 nm), blue (410–530 nm), green (455–610 nm), or red (575– 700 nm) wavebands (Figure 2b), flies were offered a choice between a full-spectrum stimulus consisting of all four wavebands (UV, blue, green, red) and a stimulus lacking one of these wavebands (Exps. 21–24; Table S1). The LED arrays were calibrated to produce illumination of near-equal total intensity (4×10^13^ photons/s/cm^2^) over the full 330–700 nm range across all treatments (Table S2).

Because the spectral purity of wavebands produced by filter stack stimuli was limited by the ratio of reflected to transmitted light, we custom-built ‘flashlight arrays’ which emitted substantially more light than the LED arrays (Table S2). Each flashlight array consisted of a cardboard holder which accommodated four Alonefire X37 Tactical Multicolor LED flashlights (Shenzhen Shiwang Technology Co. Ltd., Shengzen, GD, China) and one UV flashlight (Darkbeam SK68, Shenzhen Tianyida Electronic Technology Co. Ltd., Shengzen, GD, China) (Figures 1d,h and S1c). Each flashlight array was placed in an aluminum foil-lined plastic reflector printed by an Ender-3 Pro 3D printer (Creality, Shenzhen, GD, CN) using a black polyethylene terephthalate glycol (PETG) printing filament (1.75 mm; Amazon.com Inc., Seattle, WA, USA). This reflector was connected to a light mixing tube which, in turn, was connected to another reflector (Figure 1c,g), as described above. Using flashlight arrays, we again offered flies a choice between a full-spectrum stimulus (UV, blue, green, and red) and a stimulus lacking one of the four wavebands. However, here each paired waveband (UV, blue, green, or red) in the two stimuli, rather than all wavebands combined, had near-identical relative intensity (Exps. 25–28; Table S1).

Because the foraging activity of stable flies increases in the presence of elevated carbon dioxide (CO_2_) levels [29,47], a 30-g piece of dry ice was placed on the cage ceiling directly above, and between, the paired filter stack stimuli (Figure 1c, d) to increase landing responses of flies in experiment 21 and in all subsequent experiments.

#### H4: Stimulus intensities preferred by flies are wavelength-dependent (Exps. 29–40, S1–S12)

To determine whether fly attraction to photostimuli is mediated by the relative intensity of specific wavebands of light, flies were offered a choice between a control stimulus invariably of high relative intensity (100%) and a treatment stimulus of lower relative intensities (64–72%; 41–50% or 29–33%; Table S1), the LED pair in each set of three experiments emitting primarily UV (Exps. 29–31), blue (Exps. 32–34), green (Exps. 35–37), or red (Exps. 38–40) light (Figure 2b). It should be noted that the relative intensities of each single waveband (UV, blue, green, or red) tested in experiments 29–40 are normalized to the maximum intensity for that waveband among photostimuli used to test this hypothesis (Tables S1–S2). Further experiments tested whether stable flies can discern small relative-intensity differentials, regardless of waveband, when each of the two stimuli in a pair had low relative intensity (Exps. S1–S12; Table S1).

Stable flies, like other brachyceran flies, are spectrally insensitive to wavelengths >650 nm [12,22,35]. To narrow the waveband of the red photostimulus further beyond 650 nm, we added six layers of red lighting gel (LEE 182; Figure 1i) to the filter stack stimuli in experiments 38–40 and S10–S12.

### Statistical analyses

Data were analyzed with R statistical software (4.5.0, R Core Team 2025) using RStudio (2025.05.0+496, RStudio Team 2025). To determine the flies’ preferences for specific filter stack stimuli within each experiment, we used generalized linear models with a binomial distribution and a logit link function to compare an intercept-only model against a null model with a likelihood ratio test (LRT). To control for overdispersion, we used binomial generalized linear models with quasibinomial errors. In each experiment, we used an LRT to determine significant differences (*P* < 0.05) in the proportion of fly landing responses on each one of two paired filter stacks. Similar analyses were performed within experiments to test for differences in response preferences between male and female stable flies. Because female and male stable flies generally responded similarly in bioassays, their responses in each experiment were pooled for statistical analyses (Table S2). To compare the flies’ response preferences between experiments, we used similar generalized linear models with data from multiple experiments, with an individual intercept for each experiment. These intercepts were compared with a series of planned (*a priori*) contrasts.

## Results

### H1: Photostimuli of low relative intensity are most attractive to flies (Exps. 1–10)

Stable flies consistently preferred lower-intensity stimuli (Exps. 1–10; Figure 3), with the total number of landing responses increasing as the light intensity of photostimuli decreased (Table S3). Flies discriminated between paired low-intensity stimuli even when their relative intensity differed by as little as 2% (Exp. 10). However, the strength of the proportional preference for a lower-intensity stimulus did not differ between experiments when the relative intensity differential of stimuli was ≤8% (Exps. 8–10).

**Figure 3.**
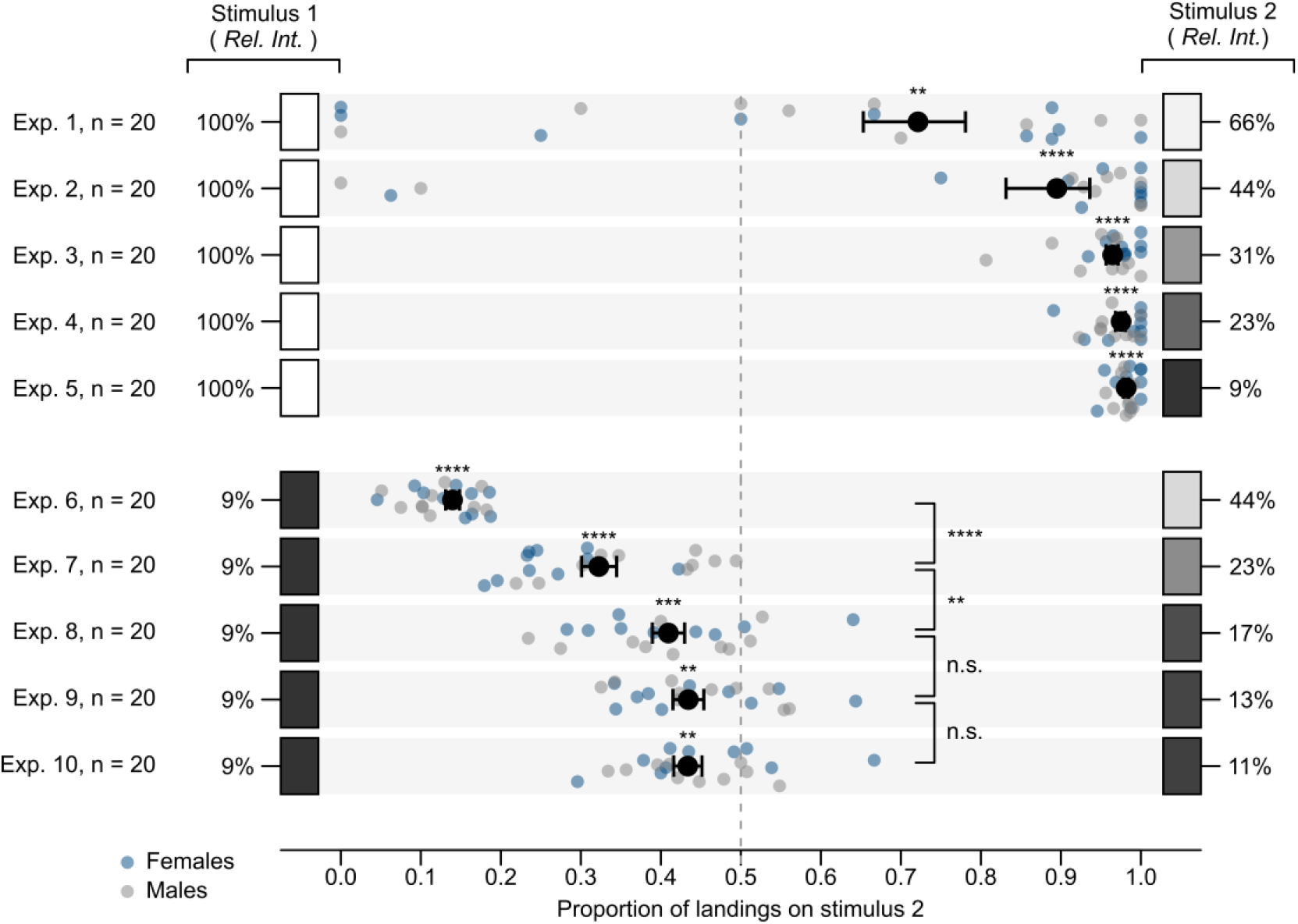
Effect of relative intensity (*Rel. Int*.) of paired photostimuli on landing responses of stable flies. The relative intensity value of each stimulus represents the mean of all wavebands transmitted through filter stacks (Table S2). Small dots show the data of individual replicates and large black dots the mean (± SEM). Asterisks (*) denote significant differences in landing responses on stimuli within and between experiments (**** *p* < 0.0001, *** *p* < 0.001, ** *p* < 0.01; ‘n. s.’ = not significant, *p* > 0.05; likelihood ratio tests).

### H2: Photostimulus intensity affects fly responses more strongly than polarization composition (Exps. 11–20)

When the relative intensity of paired photostimuli was high (≥ 58%; Exps. 11–12), flies preferentially landed on stimuli transmitting light of horizontal rather than vertical polarization (Figure 4). However, when the relative intensity of paired photostimuli was comparatively low (≤ 46%; Exps. 13–15), flies landed equally often on both stimuli, indicating that the angle of polarized light no longer affected the flies’ response preference (Figure 4).

**Figure 4.**
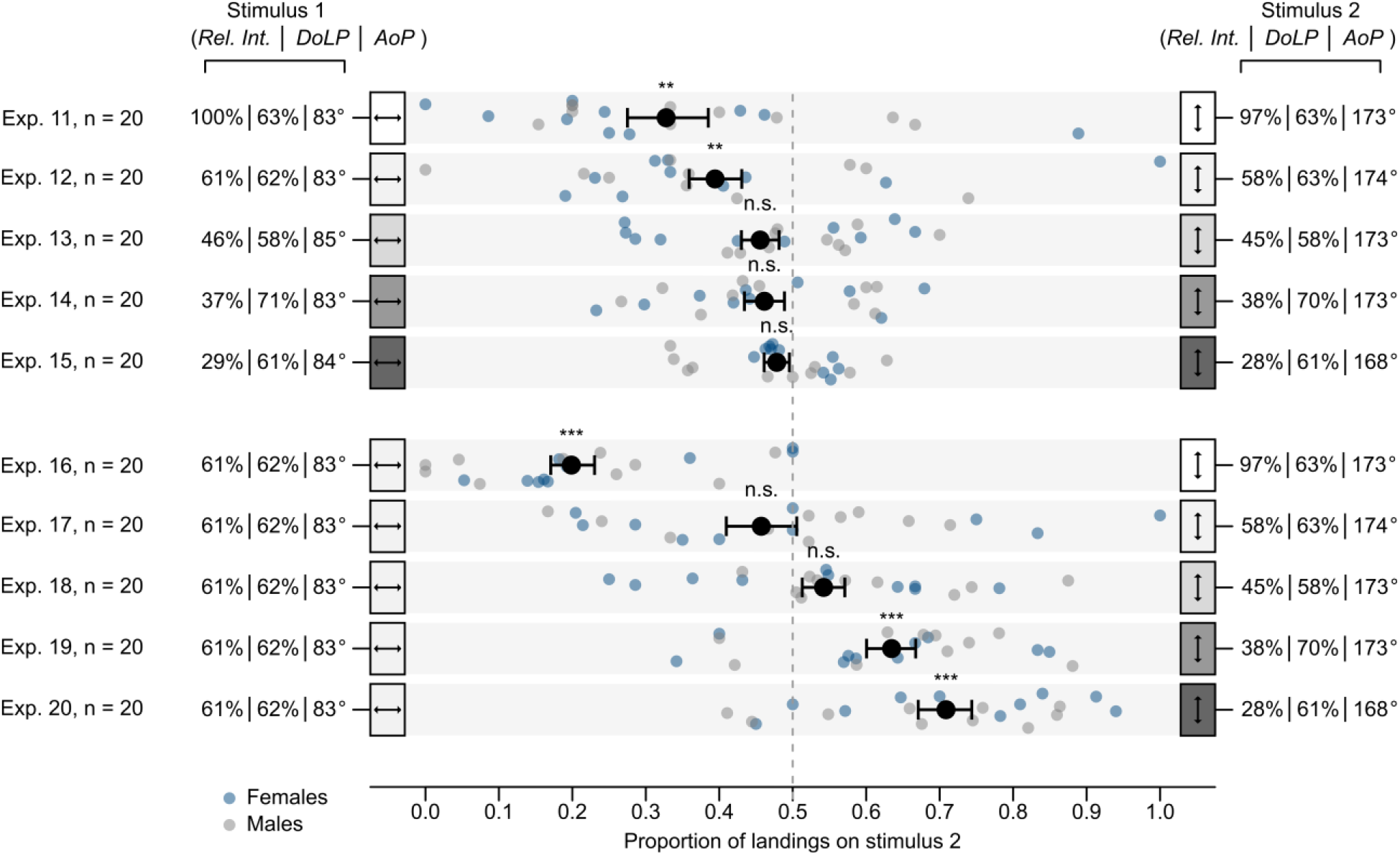
Effects of relative intensity (*Rel. Int*.) and polarization characteristics [degree of linear polarization (*DoLP*), angle of polarization (*AoP*)] of paired photostimuli on landing responses of stable flies. Flies were offered a choice between two filter stacks transmitting light of equal intensity (Exps. 11– 16) or uneven intensity (Exps. 17–20), a high *DoLP*, and an *AoP* set to predominantly be either horizontal (83–85°; stimulus 1) or vertical (168–174°; stimulus 2). The reported relative intensity, *DoLP*, and *AoP* values are the mean of all wavebands transmitted through the filter stacks (Table S2). Small dots show the data of individual replicates and large black dots the mean (± SEM). Asterisks (*) denote a significant preference for a stimulus (*** *p* < 0.001, ** *p* < 0.01; ‘n. s.’ = not significant, *p* > 0.05; likelihood ratio tests).

When flies were offered a choice between a standardized control stimulus transmitting vertically polarized light with a relative intensity of 61% and a treatment stimulus transmitting horizontally polarized light with a relative intensity of 97% (Exp. 16), 58% (Exp. 17), 45% (Exp. 18), 38% (Exp. 19) and 28% (Exp. 20), the flies’ responses shifted towards the vertically polarized stimulus as its relative intensity decreased (Figure 4). All data combined suggest that a low relative intensity of a photo stimulus has a stronger effect on stable fly attraction and landing than the angle (vertical or horizontal) of polarized light.

### H3: UV stimuli are preferentially sought by flies (Exps. 21–28)

When flies were offered a choice between paired filter stacks illuminated by either four wavebands of light (UV, blue, green, red) or any ternary combination of these wavebands, with both photostimuli in each pair set to a similar total relative intensity (81–86%), flies landed equally often on both stimuli (Exps. 21–24; Figure 5), suggesting that the omission of any waveband did not reduce the attractiveness of the stimulus.

**Figure 5.**
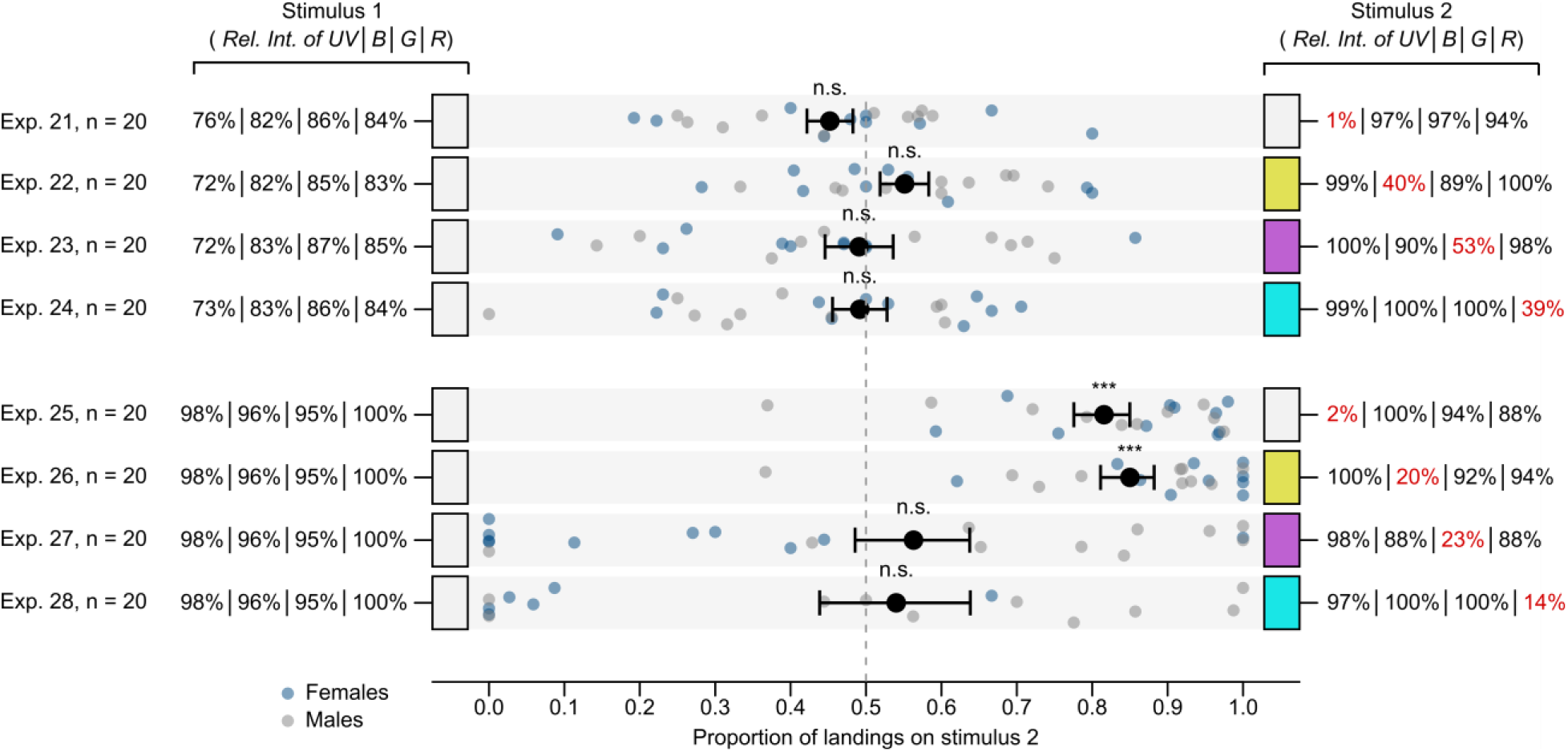
Effect of spectral composition of paired photostimuli on landing responses of stable flies. Flies were offered a choice between filter stacks that were illuminated by either a full-spectrum emission source [ultraviolet (UV; 330–395 nm), blue (B; 410–530 nm), green (G; 455–610 nm), and red (R; 575– 700 nm) wavebands (stimulus 1)], or by any ternary combination of these wavebands (stimulus 2). In experiments 21–24, paired LED arrays were calibrated to produce illumination of near-equal total intensity (4×10^13^ photons/s/cm^2^; Table S2), whereas flashlight arrays in experiments 25–28 were calibrated such that each of the paired wavebands, rather than all wavebands combined, had near-identical relative intensity. The relative intensity (*Rel. Int*.) values are reported for each waveband transmitted through the filter stacks (Table S2). Small dots show the data of individual replicates and large black symbols the mean (± SEM). Asterisks (*) denote a significant preference for stimulus 2 (*** *p* < 0.001; ‘n. s.’ = not significant, *p* > 0.05; likelihood ratio tests).

However, when flies were offered a choice between paired filter stacks illuminated by four wavebands (UV, blue, green, red) or any ternary combination of these wavebands, with each paired waveband – rather than all wavebands combined (as in experiments 21–24) – set to near-identical relative intensity (Table S2), flies landed more often on the lower-intensity (71–78%) 3-waveband stimulus than on the higher-intensity (97%) 4-waveband stimulus, provided the omitted waveband was UV (Exp. 25) or blue (Exp. 26) rather than green (Exp. 27) or red (Exp. 28) (Figure 5).

### H4: Stimulus intensities preferred by flies are wavelength-dependent (Exps. 29–40, S1–S12)

When flies were offered a choice between paired identical wavebands (UV, blue, green or red), one of which set to a high intensity (100%) and the other to a lower intensity (29–72%), flies landed more often on the lower-intensity waveband (Exps. 29–34, 36–40; Figure 6), with one exception being that flies landed equally often on green stimuli set to a relative intensity of 100% and 68% (Exp. 35; Figure 6).

**Figure 6.**
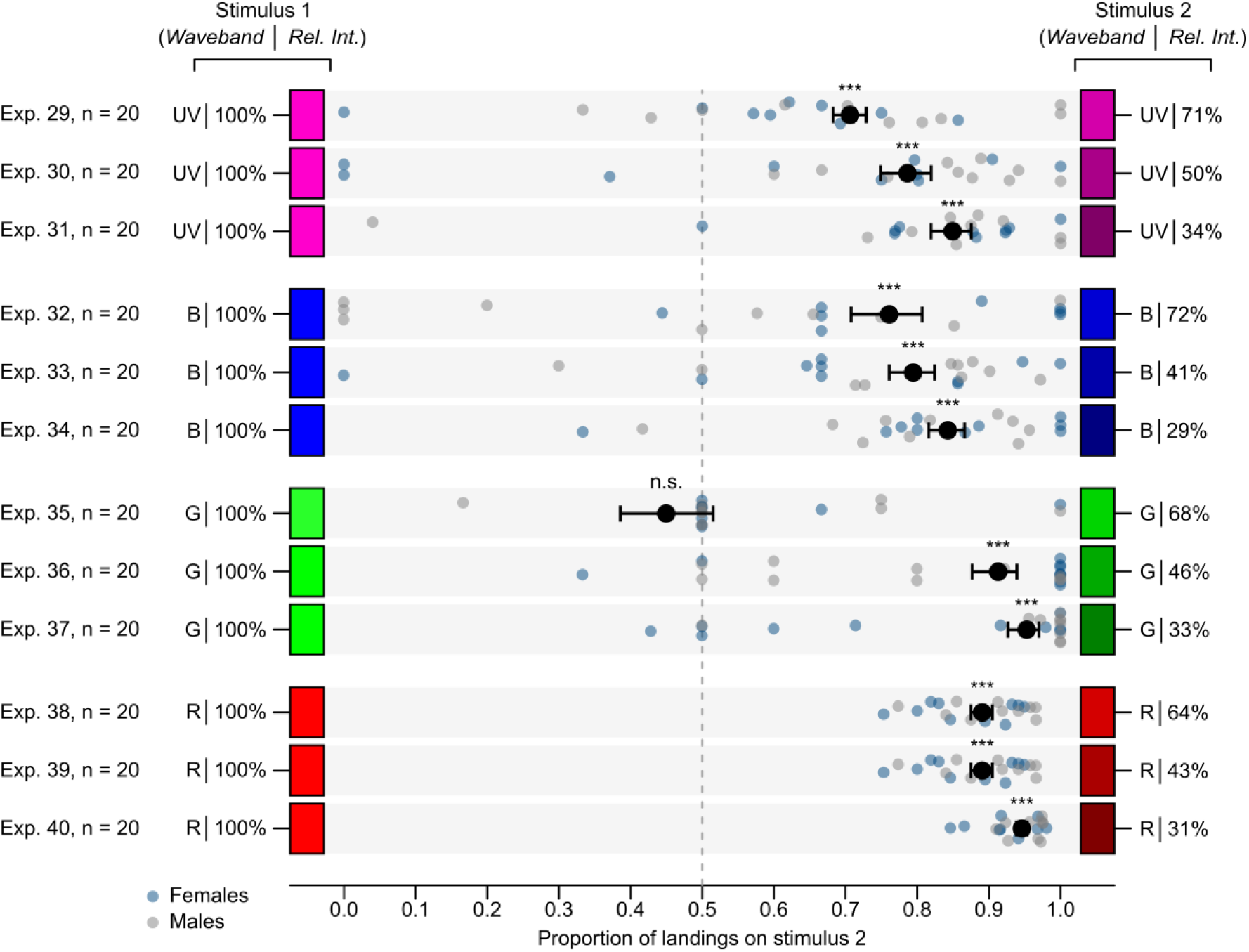
Effect of relative intensity (*Rel. Int*.) and spectral composition of paired photostimuli on landing responses of stable flies. Flies were offered a choice between filter stacks that were illuminated by an ultraviolet (UV; 330–395 nm), blue (B; 410–530 nm), green (G; 455–610 nm), or red (R; 575–700 nm) waveband of light, each waveband transmitted at either a high relative intensity (stimulus 1) or a lower relative intensity (stimulus 2) (Table S2). Small dots show the data of individual replicates and large black symbols the mean (± SEM). Asterisks (*) denote a significant preference for stimulus 2 (*** *p* < 0.001; ‘n. s.’ = not significant, *p* > 0.05; likelihood ratio tests).

When flies were offered a choice between paired identical wavebands (UV, blue, green, or red), all of which were of low relative intensity, with only modest intensity differentials between the stimuli in each pair (≤ 5% for UV, blue and green; ≤ 23% for red), flies generally failed to discriminate between intensity differentials for wavebands in the UV and the blue range (Exps. S1–S6), and generally preferred lower-intensity stimuli in the green and the red range (Exp. S7–S12) (Figure S1).

## Discussion

The intensity of photostimuli most strongly affected the visual preferences of stable flies in our experiments. Low(er)-intensity photostimuli consistently attracted flies (Figures 3, 4, 6), and the low intensity of photostimuli attracted flies more strongly than either the polarization (Figure 4) or the spectral composition (Figures 5, 6) of photostimuli. Contrary to our expectation, the presence or absence of a UV waveband in photostimuli did not affect the flies’ responses provided the total intensities of paired stimuli were near-identical (Figure 6). Because the effect of stimulus intensity prevailed over that of other stimulus characteristics, foraging stable flies appeared to not make landing decisions based primarily on true colour or polarization preferences.

Our findings that stable flies preferentially seek lower-intensity photostimuli (Figure 4) align with results obtained in previous studies on host-foraging dipterans. Tabanid flies [48], horn flies [49], mosquitoes [50], and stable flies [51] all prefer darker hosts to lighter hosts. However, the attraction of biting flies to dark hosts may not be due exclusively to the reflective intensity of these hosts. Dark hosts with a low reflective intensity also have elevated body temperature from solar warming [52,53], and they reflect light with higher degrees of polarization than lighter hosts [54]. Indeed, we have shown that stable flies preferentially land on host look-alike objects with host-like temperature [30] or a high degree of horizontally-polarized light [22]. In the current study, however, neither host-like thermal cues nor polarization cues were necessary to trigger preferential responses of flies to photostimuli with a low(er) relative intensity (Figure 3).

The lower-intensity stimulus in stimulus pairs attracted more flies regardless of its polarization characteristics (Figure 4). Similarly, stable flies preferred lower intensity photostimuli regardless of the wavelengths presented (Figures 5, 6), aligning with a previous study where object reflectance rather than object color predominantly determined landing preferences of yellow fever mosquitoes, *Aedes aegypti* [55]. If each visual modality is perceived through independent visual processing channels, then our findings could imply that stable flies rank stimulus intensity higher than other visual characteristics. Alternatively, because the intensity effect prevailed over that of other light properties, it seems conceivable that multiple visual modalities contribute to intensity perception. For instance, polarization-sensitive Japanese yellow swallowtail butterflies, *Papilio xuthus*, may perceive polarization as a modulation of intensity, whereby vertically-polarized light appears brighter than horizontally-polarized light [56]. Similarly, perception of stimulus intensity may be wavelength-dependent because responses to different wavelengths of light depend on an animal’s spectral sensitivity, at least at the retinal level. Thus, some colour-based preferences can ultimately be attributed to differences in perceived intensity. For example, *Drosophila* vinegar flies phototactically orient towards shorter rather than longer wavelengths of light [13]. However, this preference for shorter wavelengths cannot be interpreted as evidence for true colour vision because the effect can be reversed if the intensity of the shorter wavelengths is reduced [13]. If intensity perception in stable flies involves the integration of polarization and color information, stable fly responses in the current and previous study [22] could potentially be explained by a single intensity-based processing channel.

Alternatively, it remains possible that stable flies may independently process intensity, colour, and polarization information, with the effect of each modality contingent upon downstream interactions that are not yet known for stable flies. Phototactic responses of *Drosophila* are mediated by interactions between chromatic and achromatic processing channels that were previously thought to function separately [17,57]. Conceivably, color, intensity, and polarization perception may be linked and interdependent upon one another [45] rather than each type of perception functioning as a discrete modality. Notably, the manner in which different components of visual information are processed can differ considerably across invertebrate taxa [58–61], and may vary even between closely related congeners, as reported in swallowtail butterflies, *Papilio* spp. [45,56,62,63] and in common water fleas, *Daphnia* spp. [64]. Although horse flies and stable flies overlap in preferred host animals [65] and on-host feeding sites [66], they nonetheless process visual information differently. Horse flies do not prefer specific *AoPs* [27] and respond most strongly to highly polarized stimuli that lack UV or green wavelengths [21]. In comparison, stable flies exhibit a preference for horizontal polarization but only when test stimuli are composed of both UV and human-visible wavelengths [22]. Moreover, positive polarotaxis of horse flies is thought to be triggered by prospective hosts with distinct negative intensity contrast [21]. Here, we demonstrate that stable flies do not discriminate between differently polarized stimuli if stimuli have a low relative intensity (Figure 4). Because general visual preferences of haematophagous dipterans often overlap [6], it would be of considerable academic and applied interest to investigate whether foraging stable flies and more closely related biting flies (e.g., tsetse flies) or distantly related biting flies (e.g., mosquitoes) process visual host-finding cues similarly.

Stimulus intensity seems to affect stable fly responses more strongly than stimulus color, but true color vision [67] in stable flies has yet to be conclusively demonstrated. Interestingly, omission of the UV or the blue waveband, but not of the green or the red waveband, from the 4-waveband complex strongly increased the attractiveness of the complex (Exps. 25–28; Figure 5). Flies consistently also preferred the lower-intensity stimulus in stimulus pairs when both stimuli in a pair presented a red waveband (Figures 6, S1). Like other brachycerans, stable flies are spectrally insensitive to long (>650 nm) wavelengths [12,22,35]. That blue-or UV-adapted stable flies exhibit a spectral sensitivity peak above 600 nm [22] is likely the result of long wavelengths leaking through screening pigments rather than evidence of a red-sensitive receptor [68]. A similar phenomenon has been described in tsetse flies, where trap capture rates were predictable based on the red-light reflectance of traps [69] even though tsetse flies lack a red-sensitive photoreceptor [20].

Although stimulus intensity most strongly affected the responses of foraging stable flies in our study, colour and polarization cues may still have a higher relative valence in other visual contexts. Responses of fiddler crabs, *Afruca tangeri*, to looming visual stimuli are described by a ‘parallel-channel model’ where polarization and intensity information are processed simultaneously and independently, and behaviour is mediated by the channel that provides the most salient contrast [60]. Behavioural context may also affect how visual information is processed. For example, females of the orchid swallowtail butterfly, *Papilio aegeus*, appear to switch between colour-dependent and intensity-dependent polarization vision when they are ovipositing and foraging, respectively [56,62]. All our experiments were run in a single visual environment with food-deprived flies whose foraging activity was likely activated by CO_2_ unintentionally exhaled by experimentalists or intentionally liberated from dry ice (Exps. 29–40, S1–S12). This context could explain why our findings do not align with reports that materials with high reflectance and/or transmittance of sunlight are effective visual attractants for stable flies [e.g., 35,36]. Also, stable flies may respond to different visual cues depending upon their physiological state or ‘behavioural mode’. For example, hungry host-foraging stable flies might seek objects with negative contrast [29] while well-fed stable flies assemble on brightly sunlit objects for thermoregulation and mate-detection [39].

## Conclusion

Stimulus intensity is a key visual cue for attraction of foraging stable flies. Colour and polarisation characteristics of stimuli potentially modulate the attractive effect of stimulus intensity, or may play more significant roles in other contexts. Whether perceived stimulus intensity is a key driver for stable fly attraction irrespective of the visual environment, and whether the flies’ visual preferences differ with behavioural context, is yet to be investigated. If visual preferences of stable flies were to shift with behavioural context, then specific behavioural responses may also be dependent upon the flies’ physiological state and the contributions of other cues such as host odours [70–72]. Further research is needed to test whether, and to what extent, the flies’ responses are dependent upon (*i*) behavioural context and (*ii*) contrast between stimuli and the background visual environment.

## Acknowledgements

We thank Brian Anderson, Erin Anderson, and Nelson Dinn for facilitating fly collections, and Sharon Oliver for comments.

