## Supplementary Materials for "Photostimulus intensity, rather than colour or polarization, is the key driver for stable fly visual attraction in laboratory bioassays"

**Table S1.** List of laboratory behavioral experiments (Exps.), the number of replicates (n) run, and stimuli characteristics tested, including the waveband (R = red, 575–700 nm; G = green, 455–610 nm; B = blue, 410–530 nm; UV = ultraviolet, 330–395 nm), relative intensity (Rel. Int.), angle of polarization (AoP), and degree of linear polarization (DoLP); all experiments tested 3- to 7-day-old female or male stable flies.

| Exps. (n) | Stimulus 1 | | | | | | | Stimulus 2 | | | | | | |
|---|---|---|---|---|---|---|---|---|---|---|---|---|---|---|
| | Waveband | | | | Rel. Int.[1] | DoLP[2] | AoP[3] | Waveband | | | | Rel. Int.[1] | DoLP[2] | AoP[3] |
| | UV | B | G | R | | | | UV | B | G | R | | | |
| H1: Photostimuli of low relative intensity are most attractive to flies (Exps. 1–10) | | | | | | | | | | | | | | |
| 1 (20) | ✓ | ✓ | ✓ | ✓ | 100.0% | <20% | – | ✓ | ✓ | ✓ | ✓ | 65.8% | <20% | – |
| 2 (20) | ✓ | ✓ | ✓ | ✓ | 100.0% | <20% | – | ✓ | ✓ | ✓ | ✓ | 44.5% | <20% | – |
| 3 (20) | ✓ | ✓ | ✓ | ✓ | 100.0% | <20% | – | ✓ | ✓ | ✓ | ✓ | 30.5% | <20% | – |
| 4 (20) | ✓ | ✓ | ✓ | ✓ | 100.0% | <20% | – | ✓ | ✓ | ✓ | ✓ | 22.8% | <20% | – |
| 5 (20) | ✓ | ✓ | ✓ | ✓ | 100.0% | <20% | – | ✓ | ✓ | ✓ | ✓ | 9.3% | <20% | – |
| 6 (20) | ✓ | ✓ | ✓ | ✓ | 9.3% | <20% | – | ✓ | ✓ | ✓ | ✓ | 44.5% | <20% | – |
| 7 (20) | ✓ | ✓ | ✓ | ✓ | 9.3% | <20% | – | ✓ | ✓ | ✓ | ✓ | 22.8% | <20% | – |
| 8 (20) | ✓ | ✓ | ✓ | ✓ | 9.3% | <20% | – | ✓ | ✓ | ✓ | ✓ | 17.4% | <20% | – |
| 9 (20) | ✓ | ✓ | ✓ | ✓ | 9.3% | <20% | – | ✓ | ✓ | ✓ | ✓ | 13.3% | <20% | – |
| 10 (20) | ✓ | ✓ | ✓ | ✓ | 9.3% | <20% | – | ✓ | ✓ | ✓ | ✓ | 11.0% | <20% | – |
| H2: Photostimulus intensity affects fly responses more strongly than its polarization composition (Exps. 11–20) | | | | | | | | | | | | | | |
| 11 (20) | ✓ | ✓ | ✓ | ✓ | 100.0% | 63% | 83° | ✓ | ✓ | ✓ | ✓ | 96.7% | 63% | 173° |
| 12 (20) | ✓ | ✓ | ✓ | ✓ | 60.5% | 62% | 83° | ✓ | ✓ | ✓ | ✓ | 57.6% | 63% | 174° |
| 13 (20) | ✓ | ✓ | ✓ | ✓ | 45.6% | 58% | 85° | ✓ | ✓ | ✓ | ✓ | 45.1% | 58% | 173° |
| 14 (20) | ✓ | ✓ | ✓ | ✓ | 37.2% | 71% | 83° | ✓ | ✓ | ✓ | ✓ | 37.5% | 70% | 173° |
| 15 (20) | ✓ | ✓ | ✓ | ✓ | 29.2% | 61% | 84° | ✓ | ✓ | ✓ | ✓ | 28.1% | 61% | 168° |
| 16 (20) | ✓ | ✓ | ✓ | ✓ | 60.5% | 62% | 83° | ✓ | ✓ | ✓ | ✓ | 96.7% | 63% | 173° |
| 17 (20) | ✓ | ✓ | ✓ | ✓ | 60.5% | 62% | 83° | ✓ | ✓ | ✓ | ✓ | 57.6% | 63% | 174° |
| 18 (20) | ✓ | ✓ | ✓ | ✓ | 60.5% | 62% | 83° | ✓ | ✓ | ✓ | ✓ | 45.1% | 58% | 173° |
| 19 (20) | ✓ | ✓ | ✓ | ✓ | 60.5% | 62% | 83° | ✓ | ✓ | ✓ | ✓ | 37.5% | 70% | 173° |
| 20 (20) | ✓ | ✓ | ✓ | ✓ | 60.5% | 62% | 83° | ✓ | ✓ | ✓ | ✓ | 28.1% | 61% | 168° |
| H3: UV stimuli are preferentially sought by flies (Exps. 21–28) | | | | | | | | | | | | | | |
| 21 (20) | ✓ | ✓ | ✓ | ✓ | 81.6% | <20% | – | – | ✓ | ✓ | ✓ | 72.0% | <20% | – |
| 22 (20) | ✓ | ✓ | ✓ | ✓ | 80.5% | <20% | – | ✓ | – | ✓ | ✓ | 82.0% | <20% | – |
| 23 (20) | ✓ | ✓ | ✓ | ✓ | 81.7% | <20% | – | ✓ | ✓ | – | ✓ | 85.5% | <20% | – |
| 24 (20) | ✓ | ✓ | ✓ | ✓ | 81.5% | <20% | – | ✓ | ✓ | ✓ | – | 84.4% | <20% | – |
| 25 (20) | ✓ | ✓ | ✓ | ✓ | 97.3% | <20% | – | – | ✓ | ✓ | ✓ | 70.9% | <20% | – |
| 26 (20) | ✓ | ✓ | ✓ | ✓ | 97.3% | <20% | – | ✓ | – | ✓ | ✓ | 76.6% | <20% | – |
| 27 (20) | ✓ | ✓ | ✓ | ✓ | 97.3% | <20% | – | ✓ | ✓ | – | ✓ | 74.1% | <20% | – |
| 28 (20) | ✓ | ✓ | ✓ | ✓ | 97.3% | <20% | – | ✓ | ✓ | ✓ | – | 77.7% | <20% | – |
| H4: Stimulus intensities preferred by flies are wavelength-dependent (Exps. 29–40, S1–S12) | | | | | | | | | | | | | | |
| 29 (20) | ✓ | – | – | – | 100.0% | <20% | – | ✓ | – | – | – | 71.2% | <20% | – |
| 30 (20) | ✓ | – | – | – | 100.0% | <20% | – | ✓ | – | – | – | 50.2% | <20% | – |
| 31 (20) | ✓ | – | – | – | 100.0% | <20% | – | ✓ | – | – | – | 33.5% | <20% | – |
| 32 (20) | – | ✓ | – | – | 100.0% | <20% | – | – | ✓ | – | – | 71.6% | <20% | – |
| 33 (20) | – | ✓ | – | – | 100.0% | <20% | – | – | ✓ | – | – | 40.7% | <20% | – |
| 34 (20) | – | ✓ | – | – | 100.0% | <20% | – | – | ✓ | – | – | 29.4% | <20% | – |
| 35 (20) | – | – | ✓ | – | 100.0% | <20% | – | – | – | ✓ | – | 67.9% | <20% | – |
| 36 (20) | – | – | ✓ | – | 100.0% | <20% | – | – | – | ✓ | – | 46.3% | <20% | – |
| 37 (20) | – | – | ✓ | – | 100.0% | <20% | – | – | – | ✓ | – | 32.9% | <20% | – |
| 38 (20) | – | – | – | ✓ | 100.0% | <20% | – | – | – | – | ✓ | 64.2% | <20% | – |
| 39 (20) | – | – | – | ✓ | 100.0% | <20% | – | – | – | – | ✓ | 43.2% | <20% | – |
| 40 (20) | – | – | – | ✓ | 100.0% | <20% | – | – | – | – | ✓ | 31.4% | <20% | – |
| S1 (20) | ✓ | – | – | – | 1.3% | <20% | – | ✓ | – | – | – | 3.8% | <20% | – |
| S2 (20) | ✓ | – | – | – | 1.3% | <20% | – | ✓ | – | – | – | 2.6% | <20% | – |
| S3 (20) | ✓ | – | – | – | 1.3% | <20% | – | ✓ | – | – | – | 1.8% | <20% | – |
| S4 (20) | – | ✓ | – | – | 2.2% | <20% | – | – | ✓ | – | – | 5.3% | <20% | – |
| S5 (20) | – | ✓ | – | – | 2.2% | <20% | – | – | ✓ | – | – | 4.0% | <20% | – |
| S6 (20) | – | ✓ | – | – | 2.2% | <20% | – | – | ✓ | – | – | 2.9% | <20% | – |
| S7 (20) | – | – | ✓ | – | 2.2% | <20% | – | – | – | ✓ | – | 4.9% | <20% | – |
| S8 (20) | – | – | ✓ | – | 2.2% | <20% | – | – | – | ✓ | – | 3.7% | <20% | – |
| S9 (20) | – | – | ✓ | – | 2.2% | <20% | – | – | – | ✓ | – | 3.1% | <20% | – |
| S10 (20) | – | – | – | ✓ | 11.7%[4] | <20% | – | – | – | – | ✓ | 22.7%[4] | <20% | – |
| S11 (20) | – | – | – | ✓ | 11.7%[4] | <20% | – | – | – | – | ✓ | 17.1%[4] | <20% | – |
| S12 (20) | – | – | – | ✓ | 11.7%[4] | <20% | – | – | – | – | ✓ | 13.9%[4] | <20% | – |

[1] Relative intensities (photon flux) of photostimuli reported under H1–H3 in this table are the mean intensity of UV, blue, green, and red wavebands, each normalized to the maximum intensity of each waveband generated by a filter stack in each set of experiments that tested a particular hypothesis (see Table S2).

[2] Stimuli produced by filter stacks which did not include a UV-visible polarizer may have been slightly polarized by reflection.

[3] *AoP* was not estimated when stimulus light was not transmitted through a polarizer, resulting in a *DoLP* <20%.

[4] Adding red lighting gel decreased the absolute intensities of red stimuli. Consequently, intensity values of red stimuli were normalized to a lower maximum intensity, and lower relative intensity values are reported in experiments S1–S9 than in experiments S10–S12 (see Table S2).

**Table S2.** Relative intensity (Rel. Int.), degree of linear polarization (*DoLP*), and angle of polarization (*AoP*) of filter stacks (Figure 1e,i) tested in experiments (Exps.) 1–52 for behavioral responses of stable flies. Filter stacks varied in the number of light-diffusing layers of cheese cloth (Ch. Cl.) through which light passed before being transmitted through layer(s) of light-absorbing black screen (Bl. Scr.) and, in some experiments, a UV-visible light polarizer (Pol.; Exps. 11–20) or 3 layers of red lighting gel (Exps. 29–31) which narrowed the red waveband (Figure S1) and reduced its relative light intensity. Note that some configurations of filter stack stimuli were used in multiple experiments (see 'Exps.' Column). Filter stacks were illuminated by (*i*) a metal halide lamp whose light was reflected by a mirror (Exps. 1–20), (*ii*) a light-emitting diode (LED) array (Exps. 21–24), or (*iii*) a flashlight array (Exps. 25–52) whose light was channelled through the filter stack via a light-mixing tube and a reflector (Figure 1).

| Exps. | Filter stack illumination | Pol.[1] | Ch. Cl. | Bl. Scr. | UV (330–395 nm) | | | Blue (410–530 nm) | | | Green (455–610 nm) | | | Red (575–700 nm) | | |
|---|---|---|---|---|---|---|---|---|---|---|---|---|---|---|---|---|
| | | | | | Rel. Int. | *DoLP*[2] | *AoP*[3] | Rel. Int. | *DoLP*[2] | *AoP*[3] | Rel. Int. | *DoLP*[2] | *AoP*[3] | Rel. Int. | *DoLP*[2] | *AoP*[3] |
| **H1: Photostimuli of low relative intensity are most attractive to flies (Exps. 1–10)** Note: Relative intensities of filter stacks are in comparison to the relative intensity of a filter stack fitted with 1 layer of black screen illuminated by the metal halide lamp (Exps. 1–5). | | | | | | | | | | | | | | | | |
| **1–5[4]** | metal halide lamp | ✳ | 0 | 1 | 100.0% | <20% | – | 100.0% | <20% | – | 100.0% | <20% | – | 100.0% | <20% | – |
| **1** | metal halide lamp | ✳ | 0 | 2 | 68.8% | <20% | – | 64.9% | <20% | – | 64.7% | <20% | – | 64.6% | <20% | – |
| **2, 6** | metal halide lamp | ✳ | 0 | 3 | 47.0% | <20% | – | 43.9% | <20% | – | 43.5% | <20% | – | 43.4% | <20% | – |
| **3** | metal halide lamp | ✳ | 0 | 4 | 33.1% | <20% | – | 29.9% | <20% | – | 29.6% | <20% | – | 29.5% | <20% | – |
| **4, 7** | metal halide lamp | ✳ | 0 | 5 | 26.6% | <20% | – | 21.7% | <20% | – | 21.4% | <20% | – | 21.4% | <20% | – |
| **8** | metal halide lamp | ✳ | 0 | 7 | 19.0% | <20% | – | 17.0% | <20% | – | 16.7% | <20% | – | 16.8% | <20% | – |
| **9** | metal halide lamp | ✳ | 0 | 8 | 12.8% | <20% | – | 13.4% | <20% | – | 13.1% | <20% | – | 13.2% | <20% | – |
| **10** | metal halide lamp | ✳ | 0 | 9 | 10.3% | <20% | – | 11.4% | <20% | – | 11.1% | <20% | – | 11.2% | <20% | – |
| **5–10** | metal halide lamp | ✳ | 0 | 10 | 8.3% | <20% | – | 9.9% | <20% | – | 9.5% | <20% | – | 9.6% | <20% | – |
| **H2: Photostimulus intensity affects fly responses more strongly than its polarization composition (Exps. 11–20)** Note: Relative intensities of filter stacks are in comparison to the intensity of a filter stack fitted with 1 layer each of cheese cloth and black screen illuminated by the metal halide lamp, with the polarizer set to produce horizontally-polarized light (↔) (Exp. 11). Filter stack stimulus intensities were substantially decreased by the inclusion of a polarizer. | | | | | | | | | | | | | | | | |
| **1–5[4]** | metal halide lamp | ✳ | 0 | 1 | 7354% | <20% | – | 2132% | <20% | – | 1898% | <20% | – | 1718% | <20% | – |
| **11** | metal halide lamp | ↔ | 9[5] | 1 | 100.0% | 64% | 80° | 100.0% | 63% | 84° | 100.0% | 63% | 84° | 100.0% | 60% | 84° |
| **11, 16** | metal halide lamp | ↕ | 9[5] | 1 | 97.9% | 66% | 171° | 97.1% | 63% | 173° | 95.5% | 63% | 173° | 96.4% | 61% | 174° |
| **12, 16–20** | metal halide lamp | ↔ | 9[5] | 2 | 63.9% | 66% | 81° | 58.8% | 61% | 83° | 59.1% | 61% | 84° | 60.0% | 59% | 83° |
| **12, 17** | metal halide lamp | ↕ | 9[5] | 2 | 65.3% | 65% | 171° | 55.9% | 62% | 175° | 54.5% | 63% | 175° | 54.5% | 61% | 175° |
| **13** | metal halide lamp | ↔ | 9[5] | 3 | 51.0% | 60% | 81° | 43.9% | 57% | 86° | 44.1% | 58% | 86° | 43.4% | 56% | 86° |
| **13, 18** | metal halide lamp | ↕ | 9[5] | 3 | 52.1% | 56% | 168° | 43.9% | 59% | 174° | 42.4% | 58% | 174° | 42.1% | 57% | 174° |
| **14** | metal halide lamp | ↔ | 8 | 4 | 42.7% | 67% | 80° | 36.3% | 72% | 84° | 35.4% | 73% | 84° | 34.2% | 70% | 83° |
| **14, 19** | metal halide lamp | ↕ | 8 | 4 | 42.7% | 66% | 170° | 36.3% | 72% | 174° | 35.4% | 72% | 174° | 35.4% | 70% | 175° |
| **15** | metal halide lamp | ↔ | 8 | 5 | 34.4% | 56% | 85° | 27.9% | 62% | 83° | 27.3% | 64% | 83° | 27.3% | 61% | 83° |
| **15, 20** | metal halide lamp | ↕ | 8 | 5 | 33.3% | 59% | 155° | 26.5% | 62% | 172° | 26.1% | 63% | 172° | 26.4% | 61% | 173° |
| **H3: UV stimuli are preferentially sought by flies (Exps. 21–28)** Note: Relative intensities of filter stacks are in comparison to the maximum intensity (for each waveband) produced by filter stacks fitted with 4 layers of cheese cloth and 3 layers of black screen illuminated by a light-emitting diode (LED) array (Figure 1h). | | | | | | | | | | | | | | | | |
| **1–5[4]** | metal halide lamp | ✳ | 0 | 1 | 103.1% | <20% | – | 1585% | <20% | – | 2247% | <20% | – | 1826% | <20% | – |
| **21** | LED array[6] | ✳ | 4 | 3 | 75.5% | <20% | – | 81.6% | <20% | – | 85.9% | <20% | – | 83.6% | <20% | – |
| **21** | LED array[6] | ✳ | 4 | 3 | 0.8% | <20% | – | 97.3% | <20% | – | 96.5% | <20% | – | 93.6% | <20% | – |
| **22** | LED array[6] | ✳ | 4 | 3 | 71.5% | <20% | – | 82.1% | <20% | – | 85.0% | <20% | – | 83.4% | <20% | – |
| **22** | LED array[6] | ✳ | 4 | 3 | 98.5% | <20% | – | 40.4% | <20% | – | 89.2% | <20% | – | 100.0% | <20% | – |
| **23** | LED array[6] | ✳ | 4 | 3 | 72.3% | <20% | – | 83.3% | <20% | – | 86.6% | <20% | – | 84.6% | <20% | – |
| **23** | LED array[6] | ✳ | 4 | 3 | 100.0% | <20% | – | 90.4% | <20% | – | 53.1% | <20% | – | 98.3% | <20% | – |
| **24** | LED array[6] | ✳ | 4 | 3 | 73.3% | <20% | – | 82.9% | <20% | – | 86.2% | <20% | – | 83.8% | <20% | – |
| **24** | LED array[6] | ✳ | 4 | 3 | 99.3% | <20% | – | 100.0% | <20% | – | 100.0% | <20% | – | 38.5% | <20% | – |
| Note: Relative intensities of filter stacks are in comparison to the intensity of a filter stack fitted with 4 layers of cheese cloth and 6 layers of black screen illuminated by a flashlight LED array (Figure 1f) (Exps. 25–28; flashlight LED array 1). | | | | | | | | | | | | | | | | |
| **1–5[4]** | metal halide lamp | ✳ | 0 | 1 | 23.3% | <20% | – | 208.3% | <20% | – | 506.7% | <20% | – | 692.8% | <20% | – |
| **25–28** | flashlight array | ✳ | 4 | 6 | 97.9% | <20% | – | 96.0% | <20% | – | 95.3% | <20% | – | 100.0% | <20% | – |
| **25** | flashlight array | ✳ | 4 | 6 | 1.5% | <20% | – | 100.0% | <20% | – | 94.4% | <20% | – | 87.5% | <20% | – |
| **26** | flashlight array | ✳ | 4 | 6 | 100.0% | <20% | – | 20.4% | <20% | – | 92.0% | <20% | – | 94.0% | <20% | – |
| **27** | flashlight array | ✳ | 4 | 6 | 97.9% | <20% | – | 87.5% | <20% | – | 22.6% | <20% | – | 88.3% | <20% | – |
| **28** | flashlight array | ✳ | 4 | 6 | 97.4% | <20% | – | 99.9% | <20% | – | 100.0% | <20% | – | 13.5% | <20% | – |

| Exps. | Filter stack illumination | Pol.[1] | Ch. Cl. | Bl. Scr. | UV (330–395 nm) | | | Blue (410–530 nm) | | | Green (455–610 nm) | | | Red (575–700 nm) | | |
|---|---|---|---|---|---|---|---|---|---|---|---|---|---|---|---|---|
| | | | | | Rel. Int. | *DoLP*[2] | *AoP*[3] | Rel. Int. | *DoLP*[2] | *AoP*[3] | Rel. Int. | *DoLP*[2] | *AoP*[3] | Rel. Int. | *DoLP*[2] | *AoP*[3] |
| **H4: Stimulus intensities preferred by flies are wavelength-dependent (Exps. 29–52)** | | | | | | | | | | | | | | | | |
| Note: Relative intensities of filter stacks are in comparison to the maximum intensity (for each waveband) produced by filter stacks fitted with 4 layers of cheese cloth and 1 layer of black screen illuminated by a flashlight LED array (Figure 1f). Red light produced by the flashlight LED array was filtered through 6 layers of red lighting gel to produce a narrower red waveband. | | | | | | | | | | | | | | | | |
| **1–5**[4] | metal halide lamp | ✳ | 0 | 1 | 4.1% | <20% | – | 25.9% | <20% | – | 56.7% | <20% | – | 258.9% | <20% | – |
| **29–31** | flashlight array | ✳ | 4 | 1 | 100.0% | <20% | – | 5.0% | <20% | – | 11.2% | <20% | – | 57.3% | <20% | – |
| **29** | flashlight array | ✳ | 4 | 2 | 71.2% | <20% | – | 2.8% | <20% | – | 6.0% | <20% | – | 31.0% | <20% | – |
| **30** | flashlight array | ✳ | 4 | 3 | 50.2% | <20% | – | 1.8% | <20% | – | 3.8% | <20% | – | 18.6% | <20% | – |
| **31** | flashlight array | ✳ | 4 | 4 | 33.5% | <20% | – | 1.4% | <20% | – | 3.0% | <20% | – | 14.6% | <20% | – |
| **32–34** | flashlight array | ✳ | 4 | 1 | 0.6% | <20% | – | 100.0% | <20% | – | 32.6% | <20% | – | 4.3% | <20% | – |
| **32** | flashlight array | ✳ | 4 | 2 | 0.6% | <20% | – | 71.6% | <20% | – | 7.4% | <20% | – | 1.2% | <20% | – |
| **33** | flashlight array | ✳ | 4 | 3 | 0.4% | <20% | – | 40.7% | <20% | – | 4.1% | <20% | – | 1.9% | <20% | – |
| **34** | flashlight array | ✳ | 4 | 4 | 0.1% | <20% | – | 29.4% | <20% | – | 3.5% | <20% | – | 3.0% | <20% | – |
| **35–37** | flashlight array | ✳ | 4 | 1 | 0.6% | <20% | – | 20.2% | <20% | – | 100.0% | <20% | – | 24.0% | <20% | – |
| **35** | flashlight array | ✳ | 4 | 3 | 0.4% | <20% | – | 13.6% | <20% | – | 67.9% | <20% | – | 11.7% | <20% | – |
| **36** | flashlight array | ✳ | 4 | 5 | 0.6% | <20% | – | 9.2% | <20% | – | 46.3% | <20% | – | 7.0% | <20% | – |
| **37** | flashlight array | ✳ | 4 | 6 | 0.2% | <20% | – | 6.6% | <20% | – | 32.9% | <20% | – | 5.3% | <20% | – |
| **38–40** | flashlight array | ✳ | 4 | 1 | 0.3% | <20% | – | 2.9% | <20% | – | 5.7% | <20% | – | 100.0% | <20% | – |
| **38** | flashlight array | ✳ | 4 | 2 | 0.2% | <20% | – | 1.6% | <20% | – | 3.1% | <20% | – | 64.2% | <20% | – |
| **39** | flashlight array | ✳ | 4 | 3 | 0.1% | <20% | – | 1.1% | <20% | – | 2.0% | <20% | – | 43.2% | <20% | – |
| **40** | flashlight array | ✳ | 4 | 4 | 0.1% | <20% | – | 0.9% | <20% | – | 1.6% | <20% | – | 31.4% | <20% | – |
| **S1** | flashlight array | ✳ | 4 | 5 | 3.8% | <20% | – | 0.6% | <20% | – | 1.3% | <20% | – | 6.4% | <20% | – |
| **S2** | flashlight array | ✳ | 4 | 6 | 2.6% | <20% | – | 0.7% | <20% | – | 1.4% | <20% | – | 6.6% | <20% | – |
| **S3** | flashlight array | ✳ | 4 | 7 | 1.8% | <20% | – | 0.7% | <20% | – | 1.4% | <20% | – | 7.0% | <20% | – |
| **S1–S3** | flashlight array | ✳ | 4 | 8 | 1.3% | <20% | – | 0.7% | <20% | – | 1.4% | <20% | – | 6.8% | <20% | – |
| **S4** | flashlight array | ✳ | 4 | 8 | 0.1% | <20% | – | 5.3% | <20% | – | 1.5% | <20% | – | 4.2% | <20% | – |
| **S5** | flashlight array | ✳ | 4 | 9 | 0.1% | <20% | – | 4.0% | <20% | – | 1.5% | <20% | – | 4.8% | <20% | – |
| **S6** | flashlight array | ✳ | 4 | 10 | 0.1% | <20% | – | 2.9% | <20% | – | 1.5% | <20% | – | 5.3% | <20% | – |
| **S4–S6** | flashlight array | ✳ | 4 | 12 | 0.1% | <20% | – | 2.2% | <20% | – | 1.5% | <20% | – | 5.9% | <20% | – |
| **S7** | flashlight array | ✳ | 4 | 9 | 0.1% | <20% | – | 1.2% | <20% | – | 4.9% | <20% | – | 4.7% | <20% | – |
| **S8** | flashlight array | ✳ | 4 | 10 | 0.1% | <20% | – | 1.1% | <20% | – | 3.7% | <20% | – | 5.5% | <20% | – |
| **S9** | flashlight array | ✳ | 4 | 11 | 0.1% | <20% | – | 1.0% | <20% | – | 3.1% | <20% | – | 6.0% | <20% | – |
| **S7–S9** | flashlight array | ✳ | 4 | 12 | 0.1% | <20% | – | 0.8% | <20% | – | 2.2% | <20% | – | 6.5% | <20% | – |
| **S10** | flashlight array | ✳ | 4 | 8 | 0.1% | <20% | – | 0.8% | <20% | – | 1.4% | <20% | – | 22.7% | <20% | – |
| **S11** | flashlight array | ✳ | 4 | 9 | 0.1% | <20% | – | 0.7% | <20% | – | 1.3% | <20% | – | 17.1% | <20% | – |
| **S12** | flashlight array | ✳ | 4 | 10 | 0.1% | <20% | – | 0.7% | <20% | – | 1.3% | <20% | – | 13.9% | <20% | – |
| **S10–S12** | flashlight array | ✳ | 4 | 11 | 0.1% | <20% | – | 0.7% | <20% | – | 1.3% | <20% | – | 11.7% | <20% | – |

[1] Filter stacks included either a UV-visible polarizer set to produce horizontally (↔) or vertically (↕) polarized light, or they did not include a polarizer and instead transmitted diffuse light (✳; non-polarized).

[2] Stimuli produced by filter stacks not including a UV-visible polarizer may have been slightly polarized (<20% *DoLP*) by reflection of light from the overhead metal halide lamp.

[3] *AoP* was estimated when stimulus light was transmitted through a polarizer resulting in >20% *DoLP*.

[4] Characteristics of these identical filter stack treatments were measured only once. The measurements are reported in multiple locations of the table, each measurement normalized to the maximum relative intensity for each waveband in each data subset, to allow for comparisons between stimuli produced by filter stacks that differed in composition and source of illumination.

[5] Filter stacks which included a polarizer also had 8 layers of cheesecloth which were positioned to diffuse light before it passed through the polarizer. To achieve similar *DoLPs* across experiments 11–20, some filter stacks in experiments 11–13 and 15–20 were fitted with one additional layer of cheesecloth which partially diffused light after it had passed through the polarizer. To preserve a high(er) stimulus *DoLP*, a cover on the cage ceiling blocked direct illumination of filter stacks by the overhead metal halide lamp in experiments 11–20 (Figure 1b).

[6] The relative intensity of a filter stack stimulus illuminated by an LED array was modulated by (*i*) the number of black mesh layers in the filter stack and (*ii*) the light intensity of the programmatically-controlled LED array.

**Table S2**. Summary of descriptive and inferential statistical analyses of data obtained in behavioral experiments 1–52 (n = 20 each) with female and male stable flies.

| | Mean proportion of landings on stimulus 2 | | Planned contrast testing for sex-specific responses | | |
|---|---|---|---|---|---|
| Exp. | female | male | *F*-value | df | *p*-value |
| **1** | 0.76 | 0.68 | 0.403 | 18 | 0.534 |
| **2** | 0.88 | 0.90 | 0.045 | 18 | 0.834 |
| **3** | 0.98 | 0.95 | 2.457 | 18 | 0.134 |
| **4** | 0.98 | 0.97 | 0.224 | 18 | 0.642 |
| **5** | 0.98 | 0.98 | 0.809 | 18 | 0.380 |
| **6** | 0.15 | 0.13 | 1.477 | 18 | 0.240 |
| **7** | 0.27 | 0.38 | 7.467 | 18 | 0.014 |
| **8** | 0.41 | 0.41 | 0.008 | 18 | 0.929 |
| **9** | 0.42 | 0.46 | 0.909 | 18 | 0.353 |
| **10** | 0.44 | 0.43 | 0.078 | 18 | 0.784 |
| **11** | 0.68 | 0.64 | 0.099 | 18 | 0.757 |
| **12** | 0.62 | 0.59 | 0.260 | 18 | 0.617 |
| **13** | 0.58 | 0.52 | 1.690 | 18 | 0.210 |
| **14** | 0.55 | 0.53 | 0.126 | 18 | 0.727 |
| **15** | 0.50 | 0.55 | 2.382 | 18 | 0.140 |
| **16** | 0.20 | 0.20 | 0.002 | 18 | 0.970 |
| **17** | 0.43 | 0.47 | 0.192 | 18 | 0.667 |
| **18** | 0.51 | 0.56 | 0.679 | 18 | 0.421 |
| **19** | 0.61 | 0.66 | 0.497 | 18 | 0.490 |
| **20** | 0.73 | 0.70 | 0.220 | 18 | 0.644 |
| **21** | 0.46 | 0.45 | 0.030 | 18 | 0.865 |
| **22** | 0.50 | 0.59 | 2.235 | 18 | 0.152 |
| **23** | 0.38 | 0.56 | 4.555 | 18 | 0.047 |
| **24** | 0.54 | 0.46 | 1.276 | 18 | 0.274 |
| **25** | 0.88 | 0.79 | 1.243 | 18 | 0.280 |
| **26** | 0.88 | 0.82 | 0.793 | 18 | 0.385 |
| **27** | 0.20 | 0.75 | 13.985 | 18 | 0.002 |
| **28** | 0.09 | 0.77 | 19.532 | 18 | 0.001 |
| **29** | 0.66 | 0.76 | 5.293 | 18 | 0.034 |
| **30** | 0.75 | 0.86 | 2.808 | 18 | 0.111 |

| | | | | | |
|---|---|---|---|---|---|
| **31** | 0.87 | 0.83 | 0.466 | 18 | 0.503 |
| **32** | 0.85 | 0.66 | 4.432 | 18 | 0.051 |
| **33** | 0.73 | 0.84 | 3.383 | 18 | 0.082 |
| **34** | 0.86 | 0.83 | 0.364 | 18 | 0.554 |
| **35** | 0.80 | 0.40 | 1.897 | 18 | 0.211 |
| **36** | 0.96 | 0.88 | 1.011 | 18 | 0.330 |
| **37** | 0.94 | 0.97 | 0.884 | 18 | 0.362 |
| **38** | 0.87 | 0.91 | 1.590 | 18 | 0.224 |
| **39** | 0.92 | 0.92 | 0.006 | 18 | 0.941 |
| **40** | 0.94 | 0.96 | 2.073 | 18 | 0.167 |
| **S1** | 0.53 | 0.50 | 0.262 | 18 | 0.615 |
| **S2** | 0.56 | 0.46 | 2.910 | 18 | 0.105 |
| **S3** | 0.49 | 0.62 | 7.356 | 18 | 0.014 |
| **S4** | 0.46 | 0.41 | 0.371 | 18 | 0.550 |
| **S5** | 0.40 | 0.44 | 0.203 | 18 | 0.658 |
| **S6** | 0.47 | 0.48 | 0.025 | 18 | 0.877 |
| **S7** | 0.44 | 0.39 | 0.714 | 18 | 0.409 |
| **S8** | 0.42 | 0.43 | 0.017 | 18 | 0.897 |
| **S9** | 0.45 | 0.49 | 0.635 | 18 | 0.436 |
| **S10** | 0.41 | 0.46 | 1.094 | 18 | 0.310 |
| **S11** | 0.41 | 0.40 | 0.008 | 18 | 0.930 |
| **S12** | 0.35 | 0.43 | 9.825 | 18 | 0.006 |

**Table S3**. Summary of statistical data obtained by testing for significant differences (likelihood ratio test; $p < 0.05$) in the proportion of flies responding to stimuli in behavioural experiments 1–40 and S1–S12.

| | Stimulus 1 | | Stimulus 2 | | Likelihood ratio testing | | |
|---|---|---|---|---|---|---|---|
| Exp. | mean ± SE | mean prop. of landings | mean ± SE | mean prop. of landings | *F*-value | df | *p*-value |
| **1** | 7 ± 2 | 0.70 | 3 ± 1 | 0.30 | 9.93 | 1 | 0.006 |
| **2** | 27 ± 6 | 0.90 | 3 ± 1 | 0.10 | 25.49 | 1 | < 0.001 |
| **3** | 70 ± 9 | 0.96 | 3 ± 0 | 0.04 | 655.85 | 1 | < 0.001 |
| **4** | 79 ± 7 | 0.98 | 2 ± 0 | 0.02 | 716.62 | 1 | < 0.001 |
| **5** | 142 ± 14 | 0.98 | 3 ± 1 | 0.02 | 2003.40 | 1 | < 0.001 |
| **6** | 204 ± 21 | 0.86 | 33 ± 5 | 0.14 | 913.35 | 1 | < 0.001 |
| **7** | 186 ± 21 | 0.68 | 88 ± 13 | 0.32 | 58.81 | 1 | < 0.001 |
| **8** | 143 ± 15 | 0.41 | 99 ± 12 | 0.59 | 19.74 | 1 | < 0.001 |
| **9** | 176 ± 20 | 0.57 | 135 ± 13 | 0.43 | 11.41 | 1 | 0.003 |
| **10** | 174 ± 26 | 0.56 | 134 ± 20 | 0.44 | 13.90 | 1 | 0.001 |
| **11** | 7 ± 2 | 0.33 | 14 ± 4 | 0.67 | 8.78 | 1 | 0.008 |
| **12** | 15 ± 3 | 0.39 | 23 ± 4 | 0.61 | 8.37 | 1 | 0.009 |
| **13** | 22 ± 3 | 0.46 | 26 ± 5 | 0.54 | 2.97 | 1 | 0.101 |
| **14** | 31 ± 3 | 0.46 | 36 ± 4 | 0.54 | 1.99 | 1 | 0.174 |
| **15** | 35 ± 3 | 0.48 | 38 ± 4 | 0.52 | 1.60 | 1 | 0.221 |
| **16** | 19 ± 3 | 0.79 | 5 ± 1 | 0.21 | 69.74 | 1 | < 0.001 |
| **17** | 13 ± 2 | 0.54 | 11 ± 2 | 0.46 | 0.79 | 1 | 0.386 |
| **18** | 18 ± 3 | 0.46 | 21 ± 3 | 0.54 | 2.09 | 1 | 0.164 |
| **19** | 21 ± 3 | 0.36 | 37 ± 4 | 0.64 | 15.18 | 1 | < 0.001 |
| **20** | 12 ± 2 | 0.29 | 30 ± 4 | 0.71 | 27.83 | 1 | < 0.001 |

| | | | | | | | |
|---|---|---|---|---|---|---|---|
| **21** | 19 ± 4 | 0.54 | 16 ± 3 | 0.46 | 2.45 | 1 | 0.134 |
| **22** | 12 ± 2 | 0.44 | 15 ± 2 | 0.56 | 2.49 | 1 | 0.131 |
| **23** | 11 ± 2 | 0.50 | 11 ± 3 | 0.50 | 0.04 | 1 | 0.841 |
| **24** | 9 ± 1 | 0.50 | 9 ± 1 | 0.50 | 0.05 | 1 | 0.822 |
| **25** | 11 ± 3 | 0.19 | 48 ± 7 | 0.81 | 47.15 | 1 | < 0.001 |
| **26** | 9 ± 2 | 0.16 | 48 ± 7 | 0.84 | 54.97 | 1 | < 0.001 |
| **27** | 7 ± 2 | 0.41 | 10 ± 3 | 0.59 | 0.67 | 1 | 0.424 |
| **28** | 8 ± 2 | 0.47 | 9 ± 4 | 0.53 | 0.16 | 1 | 0.699 |
| **29** | 10 ± 3 | 0.30 | 23 ± 9 | 0.70 | 67.97 | 1 | < 0.001 |
| **30** | 7 ± 2 | 0.21 | 26 ± 8 | 0.79 | 47.65 | 1 | < 0.001 |
| **31** | 11 ± 3 | 0.15 | 63 ± 18 | 0.85 | 87.69 | 1 | < 0.001 |
| **32** | 3 ± 1 | 0.73 | 8 ± 3 | 0.27 | 21.14 | 1 | < 0.001 |
| **33** | 6 ± 2 | 0.80 | 24 ± 6 | 0.20 | 60.09 | 1 | < 0.001 |
| **34** | 6 ± 1 | 0.83 | 29 ± 9 | 0.17 | 108.87 | 1 | < 0.001 |
| **35** | 1 ± 1 | 0.50 | 1 ± 0 | 0.50 | 0.25 | 1 | 0.634 |
| **36** | 1 ± 0 | 0.10 | 9 ± 4 | 0.90 | 59.61 | 1 | < 0.001 |
| **37** | 1 ± 0 | 0.05 | 18 ± 8 | 0.95 | 85.27 | 1 | < 0.001 |
| **38** | 13 ± 2 | 0.11 | 108 ± 15 | 0.89 | 303.43 | 1 | < 0.001 |
| **39** | 14 ± 2 | 0.08 | 163 ± 20 | 0.92 | 373.86 | 1 | < 0.001 |
| **40** | 10 ± 2 | 0.05 | 176 ± 19 | 0.95 | 918.04 | 1 | < 0.001 |
| **S1** | 86 ± 13 | 0.48 | 92 ±10 | 0.52 | 0.25 | 1 | 0.622 |
| **S2** | 82 ± 9 | 0.50 | 82 ± 13 | 0.50 | 0.00 | 1 | 0.985 |
| **S3** | 85 ± 12 | 0.45 | 104 ± 14 | 0.55 | 3.53 | 1 | 0.076 |
| **S4** | 83 ± 13 | 0.43 | 63 ± 11 | 0.57 | 2.96 | 1 | 0.102 |
| **S5** | 75 ± 11 | 0.42 | 55 ± 8 | 0.58 | 4.39 | 1 | 0.049 |

| | | | | | | | |
|---|---|---|---|---|---|---|---|
| **S6** | 85 ± 12 | 0.48 | 77 ± 14 | 0.52 | 0.45 | 1 | 0.509 |
| **S7** | 104 ± 16 | 0.59 | 72 ± 11 | 0.41 | 7.93 | 1 | 0.011 |
| **S8** | 107 ± 15 | 0.58 | 78 ± 9 | 0.42 | 5.47 | 1 | 0.030 |
| **S9** | 92 ± 9 | 0.53 | 82 ± 9 | 0.47 | 1.55 | 1 | 0.228 |
| **S10** | 124 ± 11 | 0.56 | 96 ± 10 | 0.44 | 7.42 | 1 | 0.013 |
| **S11** | 125 ± 12 | 0.60 | 85 ± 9 | 0.40 | 15.54 | 1 | < 0.001 |
| **S12** | 148 ± 16 | 0.60 | 92 ± 12 | 0.40 | 41.45 | 1 | < 0.001 |

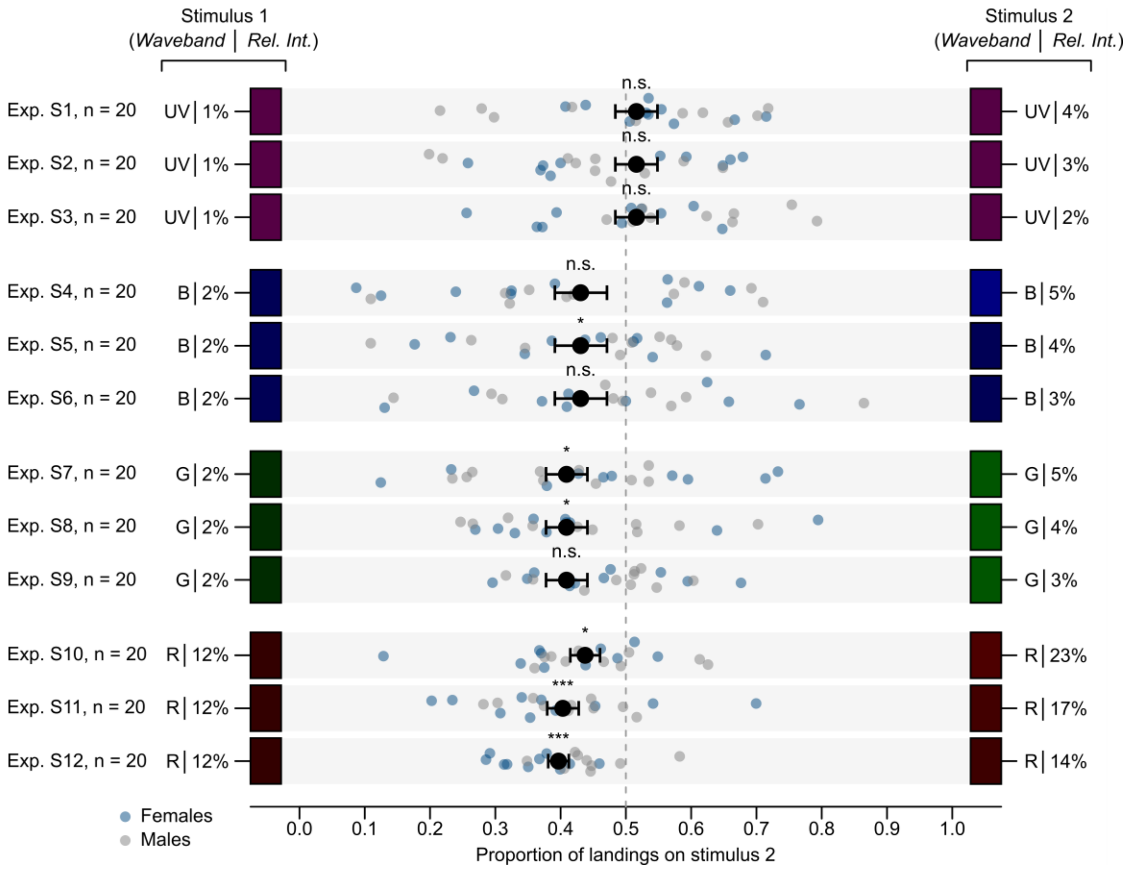


**Figure S1**. Effect of relative intensity (*Rel. Int.*) and spectral composition of paired photostimuli on landing responses of stable flies. Flies were offered a choice between filter stacks illuminated by an ultraviolet (UV; 330–395 nm), blue (B; 410–530 nm), green (G; 455–610 nm), or red (R; 575–700 nm) waveband of light, each waveband transmitting either a lower-intensity stimulus (Stimulus 1) or a higher-intensity stimulus (Stimulus 2) (Table S2). Small dots show the data of individual replicates and large black dots the mean (± SEM). Asterisks (*) denote a significant preference for stimulus 1 (*** $p < 0.001$, * $p < 0.05$; 'n. s.' = not significant, $p > 0.05$; likelihood ratio tests).
